# Gene Expression Changes in Therapeutic Ultrasound-Treated Human Chronic Wound Tissue

**DOI:** 10.1101/2022.04.13.488030

**Authors:** Olivia Boerman, Zahidur Abedin, Rose Ann DiMaria-Ghalili, Michael S. Weingarten, Michael Neidrauer, Peter A. Lewin, Kara L. Spiller

## Abstract

Low-frequency, low-intensity ultrasound has been previously shown to promote healing of chronic wounds in humans, but mechanisms behind these effects are poorly understood. The purpose of this study was to evaluate gene expression differences in debrided human venous ulcer tissue from patients treated with low-frequency (20 kHz), low-intensity (100 mW/cm^2^) ultrasound compared to a sham treatment in an effort to better understand the potential biological mechanisms. Debrided venous ulcer tissue was collected from 32 subjects one week after sham treatment or low-frequency, low-intensity ultrasound treatment. Of these samples, 7 samples (3 ultrasound treated and 4 sham treated) yielded sufficient quality total RNA for analysis by ultra-high multiplexed PCR (Ampliseq) and expression of more than 24,000 genes was analyzed. 477 genes were found to be significantly different between the ultrasound and sham groups using cut-off values of p<0.05 and fold change of 2. Gene set enrichment analysis identified 20 significantly enriched gene sets from upregulated genes and 4 significantly enriched gene sets from downregulated genes. Most of the enriched gene sets from upregulated genes were related to cellcell signaling pathways. The most significantly enriched gene set from downregulated genes was the inflammatory response gene set. These findings show that therapeutic ultrasound influences cellular behavior in chronic wounds as early as one week after application. Considering the well-known role of chronic inflammation in impairing wound healing in chronic wounds, these results suggest that a downregulation of inflammatory genes is a possible biological mechanism of ultrasound-mediated venous chronic wound healing. Such increased understanding may ultimately lead to the enhancement of ultrasound devices to accelerate chronic wound healing and increase patient quality of life.

## Introduction

Chronic venous leg ulcers (VLUs), defined as wounds that do not heal in 4-6 weeks, are a significant health care burden and affect 1-2% of the worldwide adult population with increased incidence in women and older adults [1–3]. Because of their slow healing time, VLUs cause substantial socioeconomic impact, costing up to $2 billion per year in the United States alone [4, 5]. The current standard of care for chronic VLUs include weekly wound dressing changes, the use of compression bandages, and sharp tissue debridement [5]. Although 93% of VLUs will heal in 12 months [6], the recurrence rate within 3 months is 70% [6]. There is a need for more effective chronic wound therapies.

One promising therapeutic approach is the application of low-frequency, low-intensity ultrasound (LFLI US), which has been shown in clinical trials to significantly accelerate wound healing of chronic VLUs [7–10]. In a double-blind study, patients with VLUs were treated with a nonoperational sham device or with LFLI US operating at 20 kHz and 100 mW/cm^2^ once a week for 15 minutes or 45 minutes for 8 weeks [8]. Patients in the 15-minute treatment group experienced full wound closure after 4 weeks (n=5), while patients in the sham group saw an increase in wound size (n=5), and patients in the 45-minute treatment group did not see full wound closure in the duration of the study (n=5) [8]. Despite these promising clinical results, the optimal operating parameters of therapeutic ultrasound are still unknown, and the mechanisms are poorly understood. Furthermore, these mechanisms depend on ultrasound exposure parameters, such as frequency, spatial distribution of the pressure amplitude or intensity, and time duration. A better understanding of the mechanisms of action by which therapeutic ultrasound promotes wound closure could lead to optimization of ultrasound device parameters and increase the rate of wound healing.

Researchers have explored the potential biological mechanism of LFLI US in animal models [9, 11]. However, wound healing in animals, particularly mice, is fundamentally different than wound healing in humans as it primarily occurs via contraction [12, 13] and differences in the inflammatory response to injury [14]. Therefore, there is a need to investigate mechanisms of wound healing in humans.

The goal of this study was to explore the possible hypotheses for the biological mechanisms of therapeutic ultrasound-assisted wound healing in human patients by analyzing gene expression in tissue collected from VLUs. Expression of a panel of genes has been previously linked to healing in human diabetic foot ulcers [15] and ultrasound treated VLUs [16] using tissue collected during routine wound debridement. Therefore, in this study, we similarly analyzed debrided wound tissue and assessed the whole transcriptome to identify possible mechanisms. We collected debrided human chronic wound tissue 1 week after treatment as part of an ongoing double-blinded study in which patients were treated with ultrasound operating at 20 kHz and 100 mW/cm^2^ (SPTP) or a nonoperational sham device. This tissue was processed using an ultra-high multiplexed PCR (Ampliseq) and expression of more than 24,000 genes were analyzed.

## Materials and Methods

### Study Design

As part of an ongoing double-blind human clinical study investigating the efficacy of therapeutic ultrasound for chronic wound healing (ClinicalTrials.gov Identifier: NCT03041844), sixty patients with chronic VLUs were recruited from Drexel University Comprehensive Wound Healing Program. The study protocol was reviewed and approved by the Drexel University Institutional Review Board (IRB).

Inclusion criteria for participants to enroll into the study were as follows: have a VLU that has been documented for at least 8 weeks without complete re-epithelialization, a VLU size of larger than 0.75 cm^2^, VLU must be present on the lower extremities non-weight-bearing areas. The exclusion criteria were as follows: VLU is secondary to any connective tissue disorder or blood dyscrasias, severe vascular insufficiency (ankle-brachial index lower than 0.75 or toe-brachial index below 0.5), active and untreated infection, acute deep venous thrombosis, cutaneous malignancy present on involved extremity, active (or past 6 months) cancer treatment, presence of both a diabetic ulcer and venous ulcer on the same extremity, known allergy to Tegaderm (a polyurethane dressing), pregnant woman, individuals younger than 18 years of age, prisoners, individuals not able to read or speak English, Spanish, or Mandarin, and adults unable to consent. Finally, patients with concomitant arterial disease was excluded by the presence of a palpable pedal pulse in the extremity on physical examination or a toe/bracial index of 0.6 or greater.

Weekly study visits took place in an outpatient wound clinic. Subjects randomized to the therapeutic ultrasound group were topically treated with 20kHz, 100 mW/cm^2^ for 15 minutes while patients in the sham group were treated with a non-operating device for 15 minutes. The ultrasound devices used in this study have been previously described in detail in Ngo. et al [10]. After completion of the ultrasound or sham treatment, the vascular surgeon performed tissue debridement using a surgical scalpel and participants received standard wound care. All tissue samples analyzed in this study were collected one week following ultrasound or sham treatment. Subjects who did not return to the clinic one week following ultrasound or sham treatment were considered lost to follow up.

Of the 48 subjects enrolled in the study, 16 subjects were lost to follow up resulting in 32 tissue samples processed (**Fig. 1**). Tissue samples were processed using the methods described in subsequent sections. Of the 32 processed samples, 16 did not yield RNA of sufficient quantity, and of those, 9 did not yield RNA of sufficient quality for library preparation. Thus, 7 tissue samples were ultimately analyzed, and these comprised 3 from the ultrasound treatment group and 4 from the sham group.

**Figure 1.**
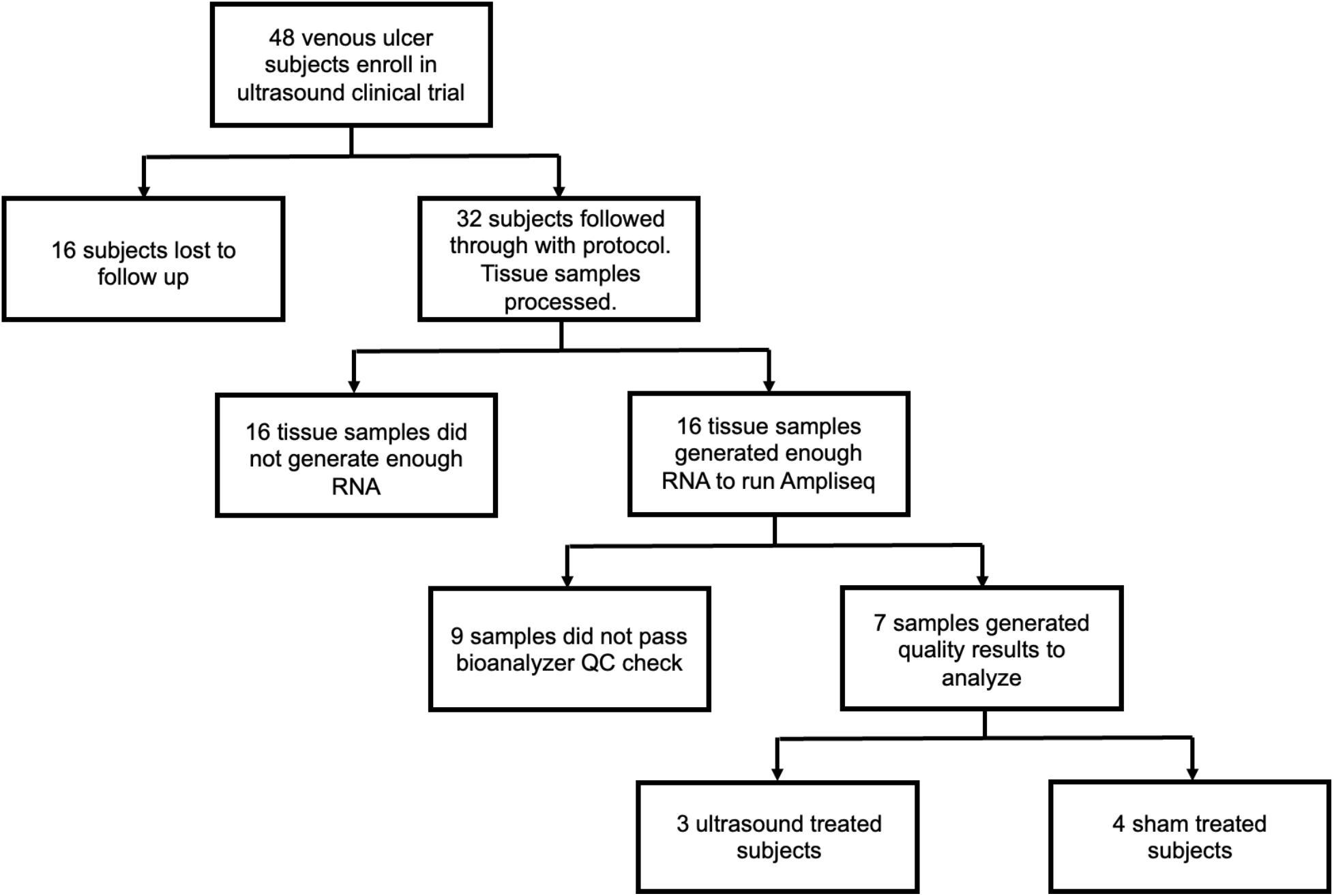
Flow chart outlining subject enrollment, sample collection, quality control, and final gene expression analysis for ultrasound-treated subjects (n=3) and sham-treated subjects (n=4).

### Debrided Tissue Collection

All participants underwent VLU debridement by a vascular surgeon as part of standard wound care regimen during each visit. Sharp debridement was conducted using a scalpel or curette after removing the overlying biofilm and necrotic tissue. Debrided tissue was immediately stored in RNAlater^®^ Solution (Ambion, Austin, TX, USA) at 4°C overnight before long term storage at −80°C prior to RNA extraction and gene expression analysis.

### RNA Extraction

The collected tissue samples were thawed at room temperature and total RNA was extracted using chloroform and Trizol method followed by purification with the Qiagen RNeasy kit (Qiagen, Inc., CA, USA) according to the manufacturer’s instructions. DNA was inactivated using DNAse I Amplification Grade (Invitrogen, Carsbad, CA, USA). Tissue samples that generated less than 10 ng of total RNA were omitted from further analysis because of insufficient RNA quantity.

### Library Preparation

cDNA libraries were constructed using Ion Ampliseq Transcriptome Human Gene Expression Kit from Thermo Fisher (MA, USA, Cat# A26325) according to manufacturer’s recommended protocol. Briefly, 10 ng of total RNA was reverse transcribed at 42°C for 30 minutes. After reverse transcription the cDNA was amplified by PCR using Ion Ampliseq Transcriptome Human Gene Expression Core Panel primers that amplified the specific targets (step 1: 99°C for 2min; step 2: 99°C for 15 sec, 60°C for 16 min for 16 cycles; step 3: Hold at 10°C). Next, the primers were partially digested as directed. Following the partial digestion of primers, adapters and bar codes were ligated to the cDNA. The cDNA was then purified using AMPure XP reagent (Beckman Coulter IN, USA, Cat#A63881) and the recommended protocol. The purified cDNA libraries were then amplified by PCR using 1X Library Amp Mix and 2 uL of 25X Library Amp Primers with the conditions as follows: Step 1: 98°C for 2 mins; Step 2: 98°C for 15 sec, 64°C for 1 min; steps 2-3 for 5 cycles and then hold at 10°C. The amplified cDNA libraries were purified using Nucleic Acid binding beads, binding buffers and processed on Agilent 2100 Bioanalyzer (Agilent CA, USA) to determine the yield and size distribution of each library.

### Bioanalyzer Library Quality Control

The quality of each final library was assessed using the Agilent^®^ dsDNA High Sensitivity Kit (Agilent CA, USA, CAT#5067-4626). A typical Ampliseq Trancriptome library is shown in Supplemental Figure 1. The molar concentration is determined and lower percentages of DNA (50 – 160 bp range) indicate a higher quality library. Those with less than 50% of the library in this region were deemed acceptable for templating and sequencing.

### Templating, Enrichment, and Sequencing

Approximately 50 pM of pooled barcoded libraries were used for templating using Thermo Fisher Ion 550 Kit-Chef (ThermoFisher, MA, USA, Cat.# A34541) and the manufacturer’s recommended protocol. Briefly, 50 pM of pooled libraries were combined and 25 ul of each sample was loaded onto the Ion Chef. Next, all reagents for the Ion 550 Chef Kit were loaded onto the Ion Chef (ThermoFisher, MA, USA) and the run was performed. The Ion Chef templated, enriched, and loaded the sample onto a 550 chip. After 15 hours the Chef paused, and QC was performed on the unenriched samples. The beads were then isolated, and quality assessment was performed on the Qubit instrument (ThermoFisher, MA, USA) to determine the percentage of beads that were polyclonal. After polyclonal assessment the Ion Chef resumed running and loaded the samples onto a 550 chip. The loaded chip was then placed into an Ion S5 sequencer, and the run was started using an Ion torrent Ampliseq transcriptome run plan that was configured based on type of library, species, number of run flows required, type of plug-in required, adapter-trimming as well as other parameters specific to the Ampliseq transcriptome run.

### Alignment and Data Analysis

After completion of the run, the raw sequence files (fastq) were aligned to the human transcriptome (hg19) reference sequences by the StrandNGS (CA, USA) software using the default parameters. The genes and transcript annotations that were used were retrieved from the Ensembl data base. Aligned BAM files were used for further analysis. Quality control was assessed by the Strand NGS program, which determined the pre- and post-alignment quality of the reads for each sample. The aligned reads were then filtered based on alignment score (≥90), match count (≤1), number of N’s allowed in reads (≤0). Aligned reads that failed these vendor quality control guidelines were removed. After this filtering, the aligned reads were normalized and quantified using the DEseq algorithm by the StrandNGS program. Next statistical analysis was performed using the Benjamini Hochberg false discovery rate and Moderated T-test to determine significant differentially expressed genes (p<0.05). After significant DEGs were identified, significant fold change between ultrasound treated subjects and sham treated subjects was determined and significant genes with a fold change >4 were clustered. Additionally, genes that had a significant fold change of 2 or higher were listed. Gene Set Enrichment Analysis (GSEA) was performed using the StrandNGS software and Broad Institute databases.

## Results

### Global gene expression analysis

A total of 7 human venous leg ulcer samples were analyzed (n=3 ultrasound; n=4 sham) for significantly different gene expression when comparing ultrasound-treated patients to sham-treated patients. 477 significantly differentially expressed genes (DEGs) were identified with a fold change greater than 2 and p<0.05 (**Fig. 2, Suppl. Tables 1 and 2**). We have also displayed the top 20 upregulated and downregulated protein encoding genes DEGs shown in **Table 1**.

**Figure 2:**
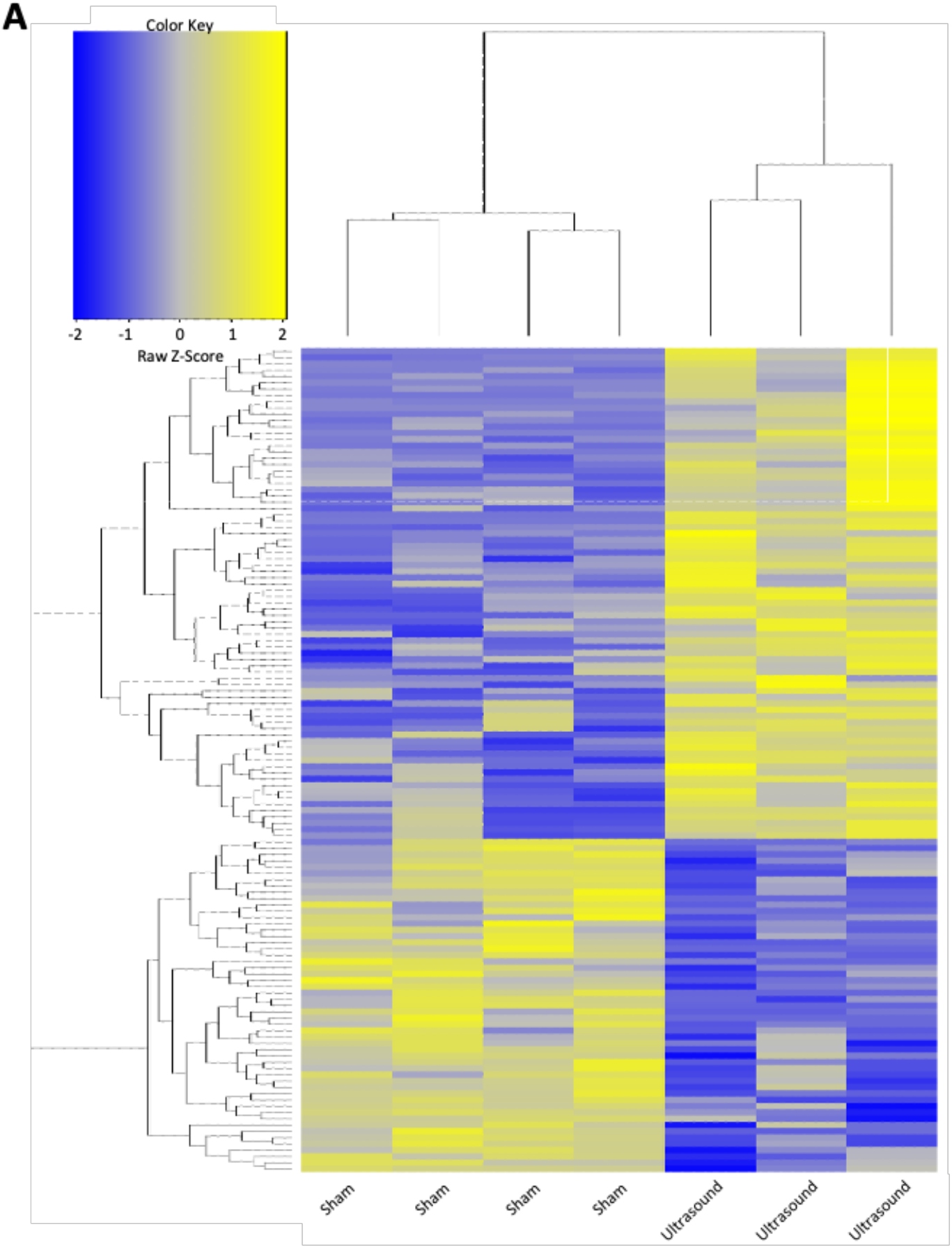
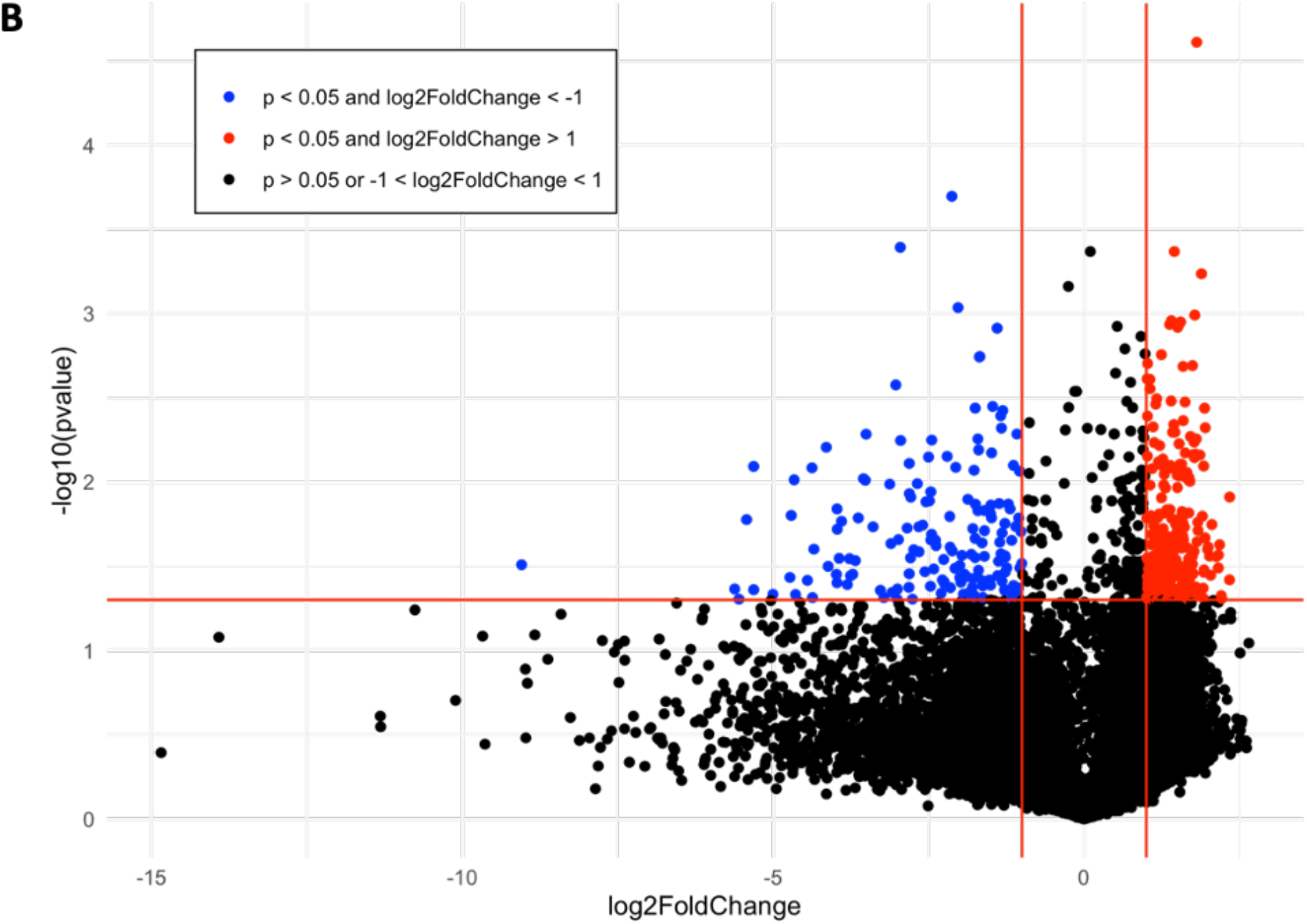
**(A)** Hierarchical clustering of significant genes (p<0.05) with a fold change >4 when comparing the ultrasound to sham group **(B)** Volcano plot of p-value vs. fold change (FC) of gene expression in ultrasound compared to sham group. Lines indicate FC greater than 2 and p<0.05 using moderated T-test with Benjamini Hochberg FDR multiple correction test.

**Table 1.** Top 20 upregulated and downregulated protein encoding DEGs (p<0.05).

| Upregulated DEGS | P-value | Fold Change | Function of Encoded Protein |
| --- | --- | --- | --- |
| PLA2G2A | 0.031 | 16.623 | Phospholipase A2 family (PLA2) which participates in the regulation of the phospholipid metabolism in biomembranes |
| MT3 | 0.041 | 12.405 | Member of the metallothionein family of genes and plays an important role in zinc and copper homeostasis, and is induced under hypoxic conditions |
| GGT3P | 0.039 | 12.222 | Involved in peptide metabolic process, response to estradiol, and response to tumor necrosis factor |
| EXTL1 | 0.043 | 11.021 | Involved in chain elongation of heparin sulfate and possibly heparan |
| RTBDN | 0.022 | 11.019 | May play a role in binding retinoids and other carotenoids as it shares homology with riboflavin binding proteins |
| CA9 | 0.043 | 10.463 | CAs are a large family of zinc metalloenzymes that catalyze the reversible hydration of carbon dioxide. May be involved in cell proliferation and transformation |
| TM4SF5 | 0.022 | 10.333 | Cell surface glycoprotein that plays a role in the regulation of cell development, activation, growth and motility |
| RORA | 0.042 | 9.297 | Member of NR1 subfamily of nuclear hormone receptors and shown to interact with NIM23-2, which is involved in organogenesis and differentiation |
| PGM3 | 0.014 | 8.882 | Member of the phosphohexose mutase family that may play a role in resistance to diabetic nephropathy and neuropathy |
| AWAT1 | 0.029 | 8.537 | Plays a central role in lipid metabolism in skin and is predominantly expressed in the sebaceous gland of the skin |
| CYP21A2 | 0.022 | 8.284 | P450 superfamily of enzymes which catalyze reactions involved in drug metabolism and synthesis of cholesterol, steroids, and other lipids |
| KIF5A | 0.003 | 7.245 | Member of the kinesin family of proteins that function as a microtubule motor in intracellular organelle transport |
| CEP290 | 0.003 | 7.101 | The presence of antibodies against this protein is associated with several forms of cancer |
| SIM2 | 0.027 | 6.994 | This gene maps within the Down syndrome chromosomal region and is thought to contribute to some specific Down syndrome phenotypes |
| USP51 | 0.028 | 6.979 | Involved in regulation of cell cycle process and enables chromatin binding activity and histone binding activity |
| CGREF1 | 0.035 | 6.793 | Predicted to enable calcium ion binding activity and predicted to be involved in negative regulation of cell proliferation |
| OR2T7 | 0.012 | 6.645 | Member of a large family of G-protein-coupled receptors and is responsible for the recognition and transduction of odorant signals |
| TMEM191C | 0.015 | 5.892 | Predicted to be an integral component of the cell membrane |
| RALA | 0.018 | 5.735 | Protein in the Ras family of proteins which mediate the transmembrane signaling initiated by the occupancy of certain cell surface receptors |
| SLC8B1 | 0.004 | 5.608 | Belongs to family of potassium-dependent sodium/calcium exchangers that maintain cellular calcium homeostasis |
| **Downregulated DEGs** | **P-value** | **Fold Change** | **Function of Encoded Protein** |
| PHACTR3 | 0.003 | -16.181 | Member of the phosphatase and actin regulator protein family and associated with the nuclear scaffold in proliferating cells |
| HGD | 0.004 | -13.439 | Enzyme homogentisate 1,2 dioxygenase and is involved in the catabolism of the amino acids tyrosine and phenylalanine |
| MED12L | 0.006 | -10.216 | Part of the Mediator complex, which is involved in transcriptional coactivation of nearly all RNA polymerase II-dependent genes |
| GP9 | 0.026 | -10.111 | A small membrane glycoprotein found on the surface of human platelets and functions as a receptor for von Willebrand factor |
| ACSM3 | 0.002 | -8.454 | Predicted to be involved in acyl-CoA metabolic process and fatty acid biosynthetic process |
| AK8 | 0.001 | -7.841 | Enables AMP binding activity; involved in nucleoside diphosphate phosphorylation and nucleoside triphosphate biosynthetic process |
| PISD | 0.01 | -7.607 | Catalyzes the conversion of phosphatidylserine to phosphatidylethanolamine in the inner mitochondrial membrane and is active in phospholipid metabolism |
| MID1 | 0.003 | -7.604 | Member of the tripartite motif (TRIM) family and likely involved in the formation of multiprotein structures acting as anchor points to microtubules |
| SIX2 | 0.011 | -7.586 | Homologous to the Drosophila 'sine oculis' homeobox protein and may be involved in limb or eye development |
| CACNA1I | 0.038 | -7.511 | Member of a subfamily of calcium channels referred to as is a low voltage-activated, T-type, calcium channel and may be involved in calcium signaling in neurons |
| IFIT1B | 0.007 | -7.433 | Predicted to act upstream of or within cellular response to interferon-alpha; cellular response to interferon-beta; and response to bacterium |
| GPA33 | 0.007 | -6.567 | Cell surface antigen that is expressed in greater than 95% of human colon cancers |
| PLB1 | 0.012 | -6.356 | Membrane-associated phospholipase that displays lysophospholipase and phospholipase A2 activities through removal of sn-1 and sn-2 fatty acids of glycerophospholipids |
| KCNQ5 | 0.007 | -6.195 | Member of the KCNQ potassium channel gene family that is differentially expressed in subregions of the brain and in skeletal muscle and Currents expressed from this protein have voltage dependences and inhibitor sensitivities in common with M-currents |
| CDC25A | 0.015 | -5.805 | Activates the cyclin-dependent kinase CDC2 by removing two phosphate groups and is required for progression from G1 to the S phase of the cell cycle |
| APOBEC3H | 0.007 | -5.78 | Cytidine deaminase that has antiretroviral activity by generating lethal hypermutations in viral genomes and are associated with increased resistance to human immunodeficiency virus type 1 infection |
| RUFY4 | 0.007 | -5.539 | Involved in autophagosome assembly; cellular response to interleukin 4; and positive regulation of macroautophagy |
| P2RY12 | 0.04 | -5.419 | Involved in platelet aggregation and is a potential target for the treatment of thromboembolisms |
| CLEC1B | 0.023 | -5.248 | Calcium-dependent (C-type) lectin-like receptors that interacts with major histocompatibility complex class I molecules and either inhibit or activate cytotoxicity and cytokine secretion |
| UBE2V1 | 0.005 | -4.986 | Involved in the control of differentiation by altering cell cycle behavior |

We employed gene set enrichment analysis to determine any significantly enriched sets of genes with similarly grouped functions, using gene sets listed on the Broad Institute database. 20 significantly enriched gene sets from upregulated DEGs were identified (**Fig. 4A**). Ten DEGs that most contributed to the enriched gene sets are shown in **Fig. 4B**. Four significantly enriched gene sets from downregulated DEGs were identified (**Fig. 5A**) with ten DEGs that most contributed to the enriched gene sets (**Fig. 5B**). Of note, the most enriched gene set from downregulated DEGs was the inflammatory response gene set, which includes 761 hallmark genes.

**Figure 4.**
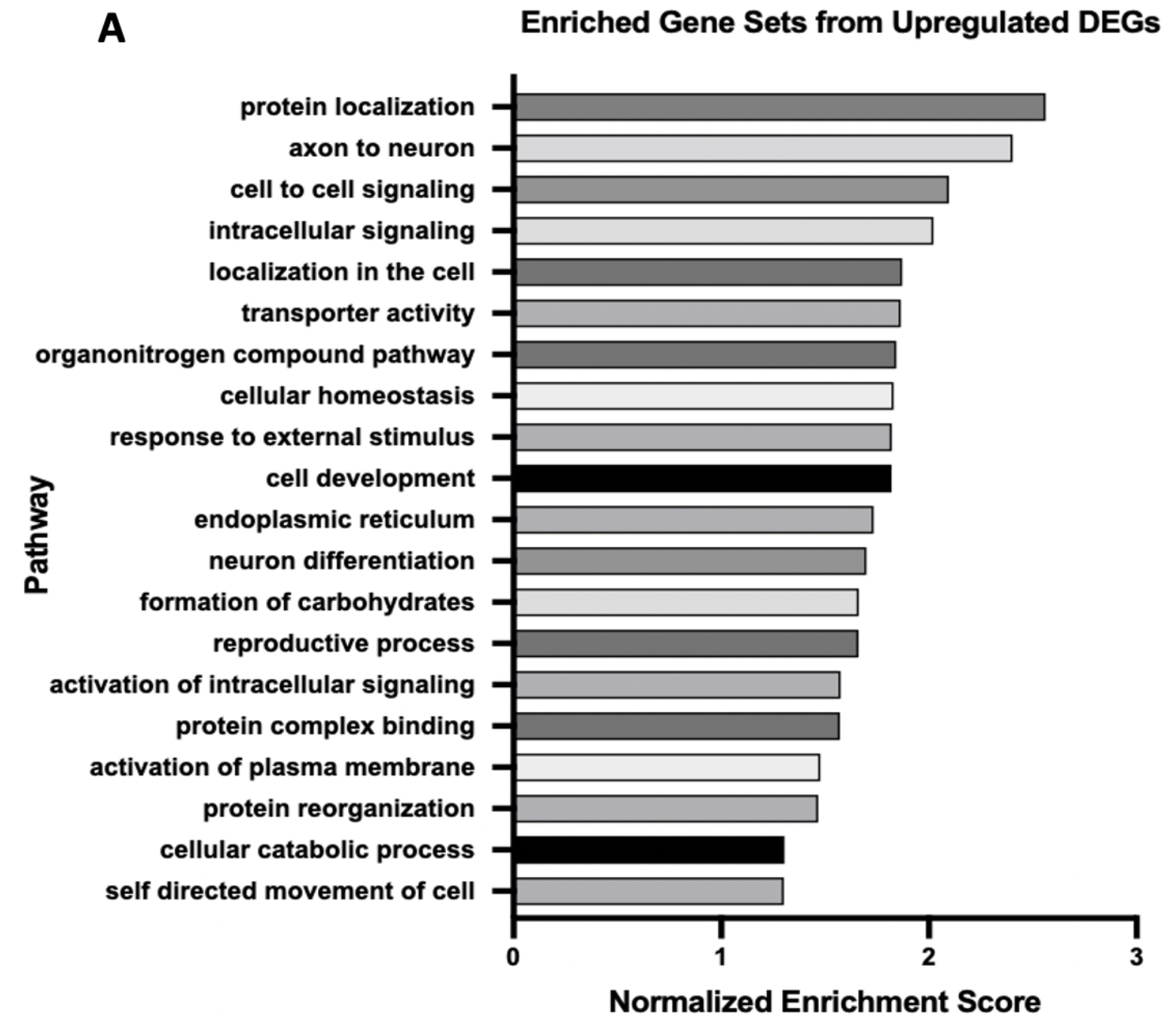
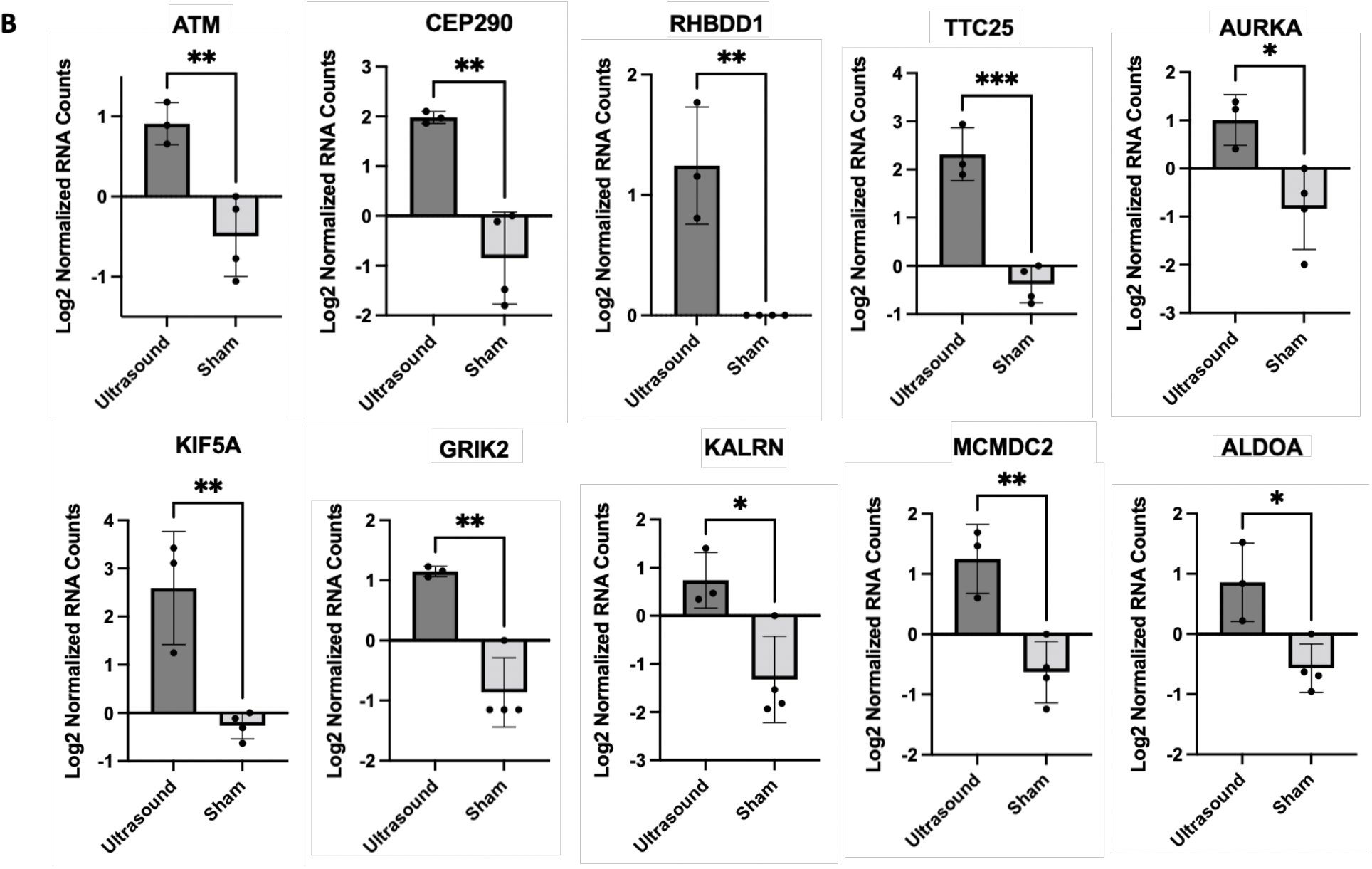
**(A)** Enriched GO terms identified from significantly upregulated DEGs (p < 0.05). **(B)** DEGs that contributed the most to the enriched gene sets; *p < 0.05; **p < 0.01, ***p < 0.001.

**Figure 5.**
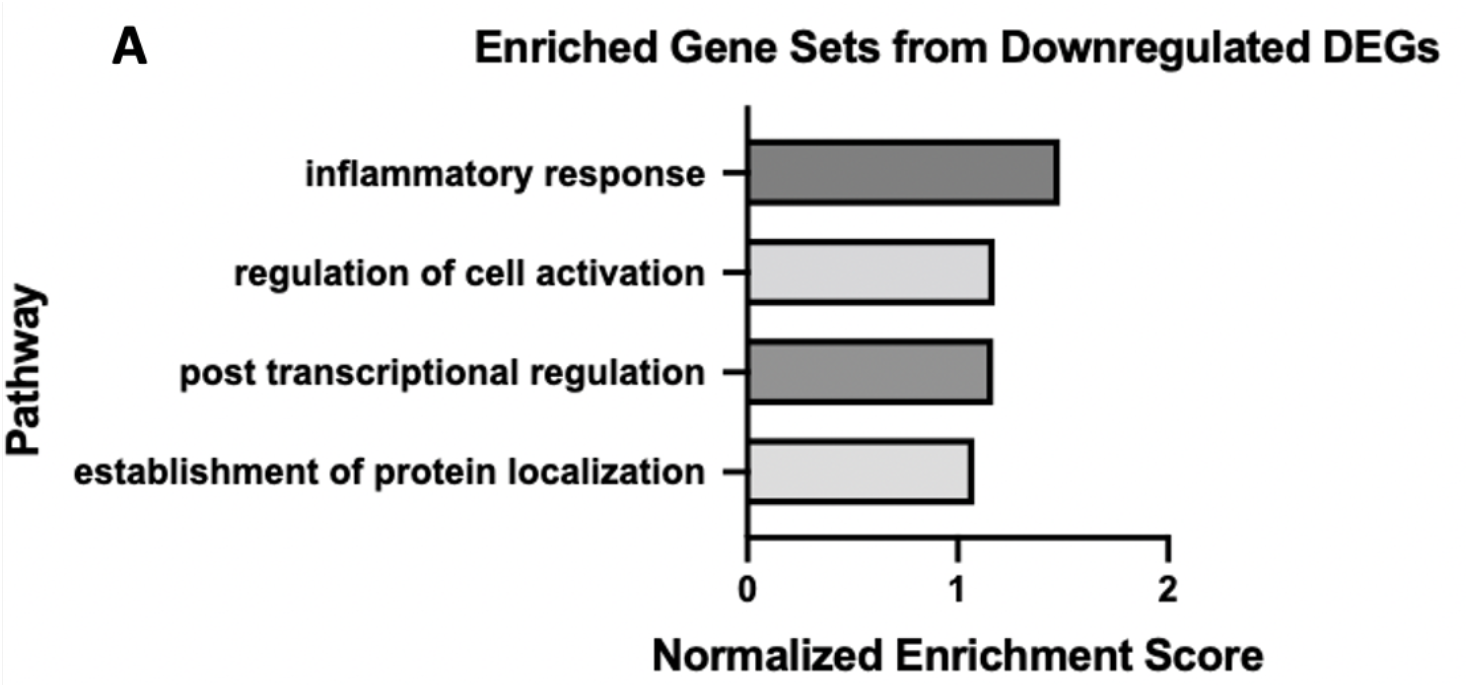
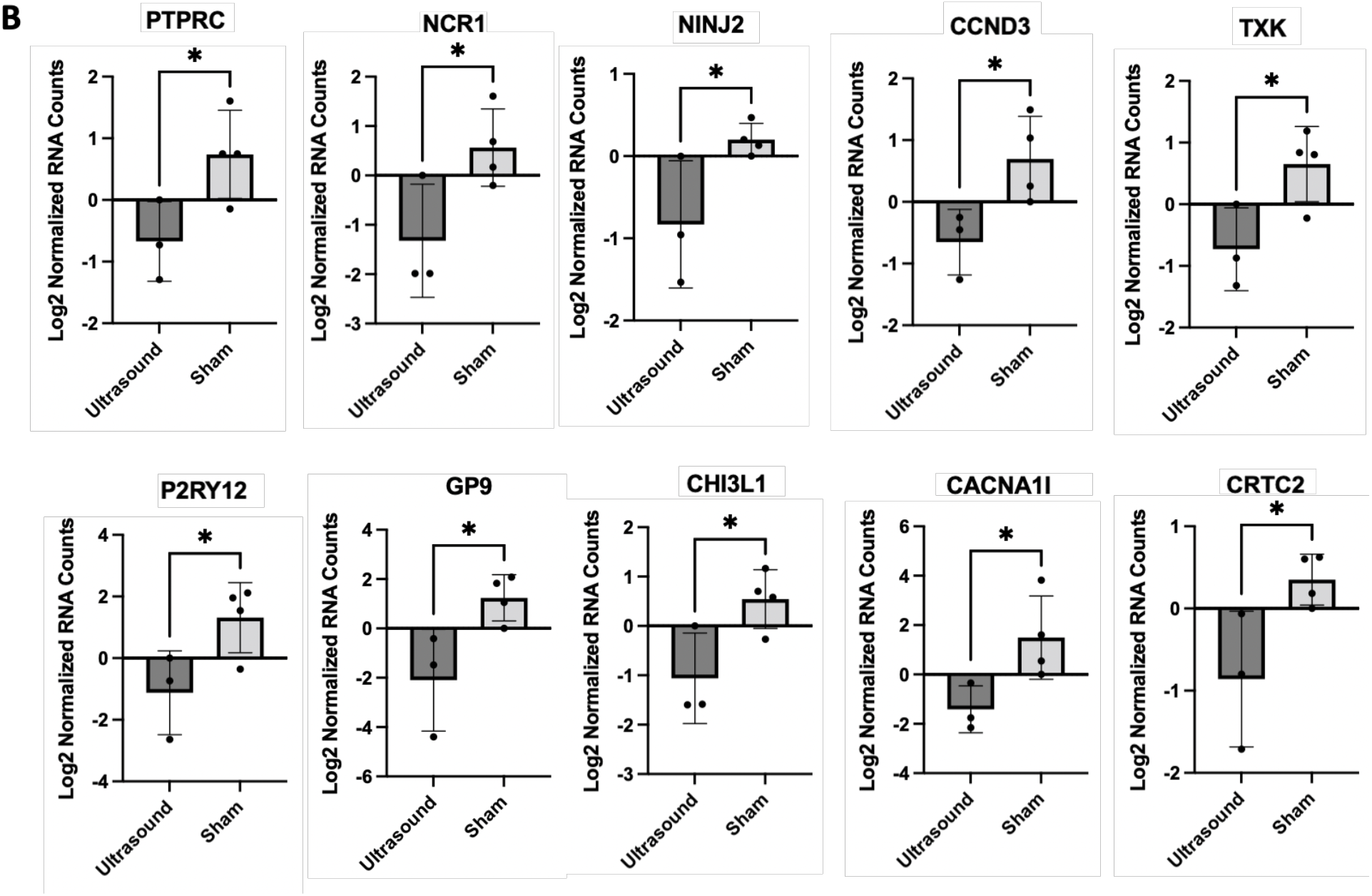
**(A)** Enriched gene sets identified from significantly downregulated DEGs (p < 0.05). **(B)** DEGs that contributed the most to the enriched gene sets; *p < 0.05.

## Discussion

Gene set enrichment analyses of human chronic venous leg wound tissue showed significant differences in participants treated with LFLI US compared to participants treated with a sham device. These results are important because they provide the foundation for future studies to investigate possible mechanisms of action behind the beneficial effects of ultrasound. In particular, the inflammatory response gene set was significantly downregulated in the ultrasound-treated patients compared to sham-treated patients as early as 1 week after treatment. These results suggest that a decrease in inflammation is a possible biological mechanism of ultrasound-mediated chronic venous leg ulcer wound healing, although this hypothesis will need to be rigorously confirmed in follow-up studies. These data contribute to the understanding of how therapeutic ultrasound modulates cells in human chronic wounds and are particularly essential in the absence of adequate animal models for human VLUs. With additional understanding of the biological mechanism of therapeutic ultrasound on cells, ultrasound parameters can be tailored to maximize chronic wound healing, ultimately leading to an increase in the quality of life for patients and a decrease in wound care associated costs.

Our findings that LFLI US significantly downregulates the inflammatory response gene set corroborates the findings of Yang et al., who found low-intensity ultrasound to globally activate anti-inflammatory genes revealed through Ingenuity Pathway Analysis (IPA) on in-vitro experimental data found in a widely available microRNA database [17]. Furthermore, Hundt et al. reported a significant downregulation in genes associated with inflammation after high-intensity (total acoustical power of 120 W) ultrasound treatment applied directly to muscle tissue in mice [18]. The downregulation of inflammation is beneficial to wound healing as sustained inflammation has been shown to be a major impediment of healing in chronic wounds [19]. In considering the literature and our findings, we suggest that the possible biological mechanisms of ultrasound-assisted healing may include both the downregulation of genes associated with inflammation and upregulation of anti-inflammatory genes. Additional studies will need to be conducted to determine specific pathways responsible for these effects. Finally, the top protein encoding genes (Table 1) can be used to motivate future studies in understanding pathway specific mechanism.

Although our findings identified several statistically significant genes and pathways that were affected by LFLI US, this study should be considered preliminary. While the study started with the recruitment of 48 venous ulcer patients, due to patient loss to follow up and poor tissue quality, the final bioinformatic analysis only included samples from 3 ultrasound and 4 sham treated subjects, thus limiting the statistical power of the study. To address these limitations, a larger cohort of patients will need to be recruited for future studies. Furthermore, this study analyzed whole tissue, which limits the understanding of individual cell types that contribute to significant differences. Hence, future work exploring cell specific mechanisms should utilize single cell analysis instead of whole tissue. Finally, we only analyzed tissue collected at a single time point (1 week after initial treatment), and other changes would be expected to occur over time. These limitations notwithstanding, these results show that low-frequency, low-intensity therapeutic ultrasound significantly affects human chronic wounds on the cellular level and these results can be used to further develop therapeutic ultrasound and maximize the potential for healing.

## Acknowledgements

This study was supported by NIH 5F31AR074847 and NIH 5R01NR015995. The content of this study is solely the responsibility of the authors and does not necessarily represent the official views of the National Institutes of Health.

## Supplemental Figures

**Supplemental Figure 1.**
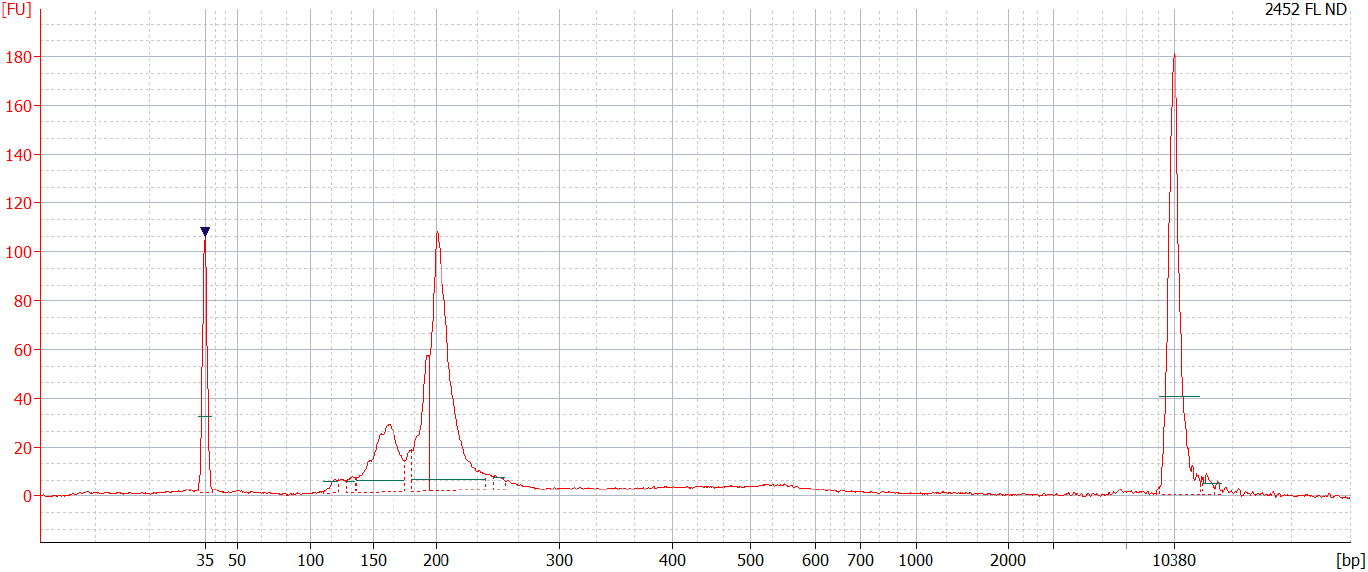
Graph displaying desired transcriptome library distribution for samples identified as optimal quality additional processing.

**Supplemental Table 1.** Table of all identified significantly upregulated genes in ultrasound group compared to sham group.

| Gene Symbol | P-value | Fold Change |
| --- | --- | --- |
| RP11-383C5.3 | 0.009 | 16.725 |
| PLA2G2A | 0.031 | 16.623 |
| MT3 | 0.041 | 12.405 |
| GGT3P | 0.039 | 12.222 |
| HIST2H2BD | 0.019 | 11.510 |
| EXTL1 | 0.043 | 11.021 |
| RTBDN | 0.022 | 11.019 |
| KRT8P46 | 0.026 | 11.013 |
| CA9 | 0.043 | 10.463 |
| TM4SF5 | 0.022 | 10.333 |
| RORA | 0.042 | 9.297 |
| PGM3 | 0.014 | 8.882 |
| AWAT1 | 0.029 | 8.537 |
| CYP21A2 | 0.022 | 8.284 |
| RP1-274L7.1 | 0.019 | 7.889 |
| AC080008.1 | 0.020 | 7.795 |
| HNRNPA3P2 | 0.046 | 7.454 |
| KIF5A | 0.003 | 7.245 |
| RP11-343H19.2 | 0.017 | 7.212 |
| CEP290 | 0.003 | 7.101 |
| SIM2 | 0.027 | 6.994 |
| CTD-2587H24.5 | 0.046 | 6.989 |
| USP51 | 0.028 | 6.979 |
| RP11-126F18.2 | 0.006 | 6.907 |
| RP4-622L5.7 | 0.041 | 6.877 |
| CGREF1 | 0.035 | 6.793 |
| OR2T7 | 0.012 | 6.645 |
| TTC25 | 0.000 | 6.489 |
| CTD-2313N18.5 | 0.005 | 6.200 |
| AP001476.2 | 0.039 | 5.969 |
| TMEM191C | 0.015 | 5.892 |
| PTPLA | 0.013 | 5.852 |
| RALA | 0.018 | 5.735 |
| LRRC46 | 0.000 | 5.678 |
| SLC8B1 | 0.004 | 5.608 |
| RP11-159J3.1 | 0.034 | 5.529 |
| RP11-302I18.1 | 0.005 | 5.484 |
| GPR39 | 0.001 | 5.398 |
| C1QTNF2 | 0.005 | 5.360 |
| ABCA9 | 0.004 | 5.307 |
| RP11-118B18.2 | 0.020 | 5.287 |
| CORIN | 0.028 | 5.180 |
| NMNAT2 | 0.005 | 5.153 |
| BPIFB1 | 0.001 | 5.090 |
| RP11-416I2.1 | 0.025 | 5.090 |
| LEMD1 | 0.022 | 5.077 |
| VSIG8 | 0.024 | 4.972 |
| PSORS1C3 | 0.044 | 4.934 |
| ZNF878 | 0.038 | 4.888 |
| RP11-461O7.1 | 0.004 | 4.822 |
| WDR93 | 0.024 | 4.796 |
| RP11-362F19.2 | 0.007 | 4.758 |
| RD3 | 0.030 | 4.732 |
| RP11-282O18.3 | 0.032 | 4.647 |
| AGBL4 | 0.007 | 4.640 |
| ELAVL2 | 0.012 | 4.619 |
| SOGA3 | 0.026 | 4.609 |
| GTF2IRD2P1 | 0.023 | 4.606 |
| C4BPB | 0.042 | 4.595 |
| CTD-2293H3.1 | 0.007 | 4.531 |
| FRG2C | 0.019 | 4.502 |
| CDK18 | 0.045 | 4.469 |
| RP3-337H4.8 | 0.006 | 4.415 |
| RP11-1250I15.3 | 0.036 | 4.399 |
| RP11-609N14.1 | 0.029 | 4.341 |
| ZNF845 | 0.040 | 4.280 |
| PPP4R4 | 0.005 | 4.246 |
| CMP21-97G8.2 | 0.038 | 4.229 |
| ZNF471 | 0.002 | 4.227 |
| AC093838.4 | 0.043 | 4.218 |
| KALRN | 0.014 | 4.172 |
| IL17B | 0.032 | 4.126 |
| EDNRA | 0.047 | 4.083 |
| RP3-499B10.3 | 0.017 | 4.082 |
| NEURL3 | 0.018 | 4.045 |
| GRIK2 | 0.001 | 4.030 |
| NUDT1 | 0.014 | 4.019 |
| TMF1 | 0.026 | 4.001 |
| AC113167.1 | 0.016 | 3.995 |
| RP11-386I23.1 | 0.007 | 3.989 |
| RP13-578N3.3 | 0.048 | 3.989 |
| SLC13A3 | 0.036 | 3.982 |
| CTD-2542L18.1 | 0.019 | 3.978 |
| RP11-564A8.4 | 0.049 | 3.963 |
| COL8A2 | 0.013 | 3.951 |
| CCL16 | 0.019 | 3.949 |
| AC017076.5 | 0.046 | 3.949 |
| ITGB1BP2 | 0.011 | 3.937 |
| RP11-53B2.2 | 0.033 | 3.927 |
| PTOV1-AS1 | 0.020 | 3.915 |
| MICALL2 | 0.012 | 3.909 |
| AC020922.1 | 0.006 | 3.892 |
| AC009499.1 | 0.013 | 3.870 |
| ANGPTL6 | 0.044 | 3.869 |
| PRSS45 | 0.001 | 3.801 |
| CTC-573M9.1 | 0.038 | 3.794 |
| RP11-394I13.2 | 0.006 | 3.739 |
| PTPN5 | 0.029 | 3.705 |
| MCMDC2 | 0.004 | 3.685 |
| RP11-169K16.9 | 0.001 | 3.683 |
| PPP1R3F | 0.007 | 3.673 |
| AP000662.4 | 0.016 | 3.659 |
| GSTO2 | 0.045 | 3.659 |
| RP11-429J17.2 | 0.039 | 3.649 |
| DHRS12 | 0.006 | 3.646 |
| WNT9A | 0.037 | 3.643 |
| AURKA | 0.017 | 3.595 |
| CYB5RL | 0.027 | 3.585 |
| PLA2G4C | 0.001 | 3.575 |
| RP11-455G16.1 | 0.035 | 3.567 |
| ITPKA | 0.047 | 3.550 |
| FRY-AS1 | 0.008 | 3.549 |
| GLULP4 | 0.019 | 3.547 |
| TIMM8A | 0.014 | 3.531 |
| PDILT | 0.007 | 3.531 |
| THUMPD1 | 0.046 | 3.499 |
| FAM131B | 0.006 | 3.494 |
| SHISA2 | 0.028 | 3.456 |
| ALMS1-IT1 | 0.023 | 3.449 |
| RP11-798M19.6 | 0.016 | 3.434 |
| SH3GL1 | 0.014 | 3.407 |
| STX1A | 0.043 | 3.394 |
| FKBPL | 0.039 | 3.387 |
| SBSPON | 0.030 | 3.372 |
| ZC3H12B | 0.019 | 3.362 |
| RP11-893F2.5 | 0.049 | 3.348 |
| RP1-212P9.2 | 0.000 | 3.321 |
| AC005086.2 | 0.017 | 3.320 |
| SCN2B | 0.003 | 3.289 |
| RP11-102M11.2 | 0.025 | 3.269 |
| RP11-227G15.9 | 0.018 | 3.258 |
| AC002310.13 | 0.047 | 3.254 |
| RP11-61A14.4 | 0.021 | 3.250 |
| ZNF232 | 0.048 | 3.246 |
| TNNI1 | 0.028 | 3.237 |
| LLPH | 0.028 | 3.228 |
| RP11-430N14.4 | 0.004 | 3.208 |
| FAM71F2 | 0.044 | 3.182 |
| NLRP8 | 0.013 | 3.147 |
| IL5RA | 0.021 | 3.145 |
| RPS15P5 | 0.022 | 3.132 |
| ABO | 0.001 | 3.126 |
| BTG3 | 0.048 | 3.120 |
| CTC-550B14.7 | 0.002 | 3.108 |
| ADCY4 | 0.046 | 3.085 |
| FABP7P2 | 0.026 | 3.078 |
| AC018865.8 | 0.027 | 3.060 |
| RP11-492E3.2 | 0.028 | 3.057 |
| RPL7P4 | 0.006 | 3.051 |
| RP11-545A16.1 | 0.006 | 3.050 |
| CA11 | 0.033 | 3.048 |
| PTGIR | 0.044 | 3.046 |
| RP5-1166F10.1 | 0.011 | 3.042 |
| CYP17A1 | 0.040 | 3.036 |
| ZNF606 | 0.029 | 3.020 |
| SFTA2 | 0.038 | 3.017 |
| MCHR2 | 0.001 | 3.017 |
| RP6-206I17.1 | 0.017 | 3.009 |
| WBP2NL | 0.026 | 3.001 |
| HS3ST3B1 | 0.041 | 3.000 |
| SMAD9 | 0.046 | 2.983 |
| CADPS | 0.047 | 2.960 |
| AICDA | 0.024 | 2.921 |
| GALNT16 | 0.043 | 2.919 |
| RNF212 | 0.020 | 2.901 |
| CICP27 | 0.041 | 2.884 |
| RP11-723D22.3 | 0.029 | 2.883 |
| RP11-283I3.6 | 0.019 | 2.875 |
| FAM173A | 0.042 | 2.866 |
| RP11-342C24.8 | 0.011 | 2.849 |
| RP11-382D8.5 | 0.016 | 2.846 |
| C6ORF165 | 0.020 | 2.843 |
| OR2H1 | 0.046 | 2.839 |
| ZNF720 | 0.036 | 2.798 |
| KB-1471A8.1 | 0.019 | 2.797 |
| ADAM29 | 0.021 | 2.788 |
| RP11-445O3.3 | 0.023 | 2.785 |
| GS1-259H13.2 | 0.008 | 2.777 |
| LINC00996 | 0.029 | 2.769 |
| PEX5L | 0.023 | 2.752 |
| RN7SL275P | 0.031 | 2.747 |
| RP11-475A13.1 | 0.049 | 2.740 |
| KLHDC1 | 0.028 | 2.735 |
| AC010970.2 | 0.007 | 2.719 |
| XXbac-B444P24.8 | 0.024 | 2.708 |
| RP11-1094M14.11 | 0.042 | 2.702 |
| WIPF3 | 0.045 | 2.702 |
| RP11-814P5.1 | 0.008 | 2.692 |
| ALDOA | 0.012 | 2.691 |
| CA14 | 0.043 | 2.669 |
| FAM157A | 0.034 | 2.657 |
| INTU | 0.015 | 2.654 |
| C9orf117 | 0.035 | 2.650 |
| ATM | 0.005 | 2.648 |
| RP11-205M3.3 | 0.043 | 2.639 |
| FBXL12 | 0.042 | 2.633 |
| TRAF5 | 0.019 | 2.629 |
| C1orf220 | 0.027 | 2.621 |
| C1ORF220 | 0.027 | 2.621 |
| LONRF2 | 0.041 | 2.601 |
| TMEM132B | 0.047 | 2.601 |
| DEF8 | 0.033 | 2.588 |
| DPH3P1 | 0.009 | 2.579 |
| RN7SL59P | 0.001 | 2.574 |
| FAM166A | 0.035 | 2.566 |
| RP11-169D4.1 | 0.021 | 2.532 |
| AL359878.1 | 0.018 | 2.521 |
| AC018506.1 | 0.019 | 2.519 |
| ING1 | 0.016 | 2.508 |
| EXOSC6 | 0.026 | 2.501 |
| RP11-356C4.3 | 0.017 | 2.496 |
| ZSCAN10 | 0.004 | 2.496 |
| RP11-521B24.3 | 0.048 | 2.493 |
| CTD-2369P2.2 | 0.012 | 2.485 |
| CORO2B | 0.032 | 2.475 |
| TCEANC2 | 0.027 | 2.474 |
| TAPBP | 0.048 | 2.470 |
| AHNAK | 0.030 | 2.469 |
| CCDC113 | 0.032 | 2.465 |
| KRTCAP2 | 0.005 | 2.451 |
| CDH17 | 0.045 | 2.434 |
| AGXT | 0.034 | 2.428 |
| LL22NC03-63E9.3 | 0.036 | 2.415 |
| RUNX1 | 0.012 | 2.410 |
| TTC16 | 0.030 | 2.395 |
| CEP112 | 0.018 | 2.390 |
| ZNF90P1 | 0.034 | 2.390 |
| RP3-368A4.5 | 0.041 | 2.388 |
| C19orf18 | 0.016 | 2.377 |
| RP5-961K14.2 | 0.023 | 2.370 |
| RHBDD1 | 0.002 | 2.369 |
| GGT6 | 0.046 | 2.364 |
| AP4M1 | 0.042 | 2.364 |
| FLJ27365 | 0.017 | 2.355 |
| PARPBP | 0.044 | 2.355 |
| DDX50P1 | 0.032 | 2.354 |
| AC003102.3 | 0.047 | 2.341 |
| BOC | 0.042 | 2.340 |
| ZNF814 | 0.025 | 2.339 |
| SPTY2D1-AS1 | 0.016 | 2.332 |
| AAED1 | 0.002 | 2.332 |
| RP11-579E24.1 | 0.021 | 2.331 |
| LNP1 | 0.042 | 2.320 |
| AC074117.10 | 0.034 | 2.313 |
| RP5-930J4.4 | 0.013 | 2.311 |
| GUSBP2 | 0.049 | 2.299 |
| AC092933.4 | 0.040 | 2.298 |
| MXRA7 | 0.033 | 2.295 |
| SPEF2 | 0.024 | 2.287 |
| VPS37A | 0.017 | 2.282 |
| DAW1 | 0.014 | 2.277 |
| CTD-2328D6.1 | 0.004 | 2.271 |
| CENPE | 0.016 | 2.266 |
| TM6SF2 | 0.014 | 2.250 |
| DZANK1 | 0.035 | 2.246 |
| RC3H1-IT1 | 0.003 | 2.229 |
| RP4-591N18.2 | 0.036 | 2.227 |
| AL161915.1 | 0.027 | 2.224 |
| HFE | 0.041 | 2.205 |
| N4BP2 | 0.037 | 2.204 |
| C1orf85 | 0.050 | 2.203 |
| RP4-535B20.4 | 0.006 | 2.190 |
| LRRC34 | 0.022 | 2.189 |
| RP11-273G15.2 | 0.016 | 2.182 |
| RBBP8 | 0.041 | 2.179 |
| RN7SL669P | 0.030 | 2.177 |
| NME9 | 0.044 | 2.176 |
| HLA-L | 0.018 | 2.164 |
| CENPQ | 0.040 | 2.157 |
| NPAP1P2 | 0.030 | 2.149 |
| CTC-338M12.6 | 0.012 | 2.144 |
| PPCS | 0.047 | 2.143 |
| PLA2G12A | 0.040 | 2.132 |
| C12orf60 | 0.044 | 2.132 |
| RP11-219G17.4 | 0.037 | 2.124 |
| AL135901.1 | 0.002 | 2.119 |
| CD109 | 0.040 | 2.118 |
| TBCCD1 | 0.008 | 2.116 |
| NAB2 | 0.030 | 2.110 |
| ITGB8 | 0.023 | 2.106 |
| USP46 | 0.025 | 2.105 |
| RNF144A | 0.025 | 2.105 |
| AC006129.2 | 0.024 | 2.092 |
| PLEKHH3 | 0.049 | 2.085 |
| POLK | 0.037 | 2.082 |
| RP11-1020A11.2 | 0.049 | 2.073 |
| DDX19A | 0.024 | 2.070 |
| NAGPA | 0.026 | 2.063 |
| RP11-84A19.4 | 0.049 | 2.062 |
| ZC3H12C | 0.035 | 2.061 |
| FASTKD5 | 0.040 | 2.056 |
| SCNN1D | 0.041 | 2.052 |
| RP11-547D24.1 | 0.032 | 2.047 |
| RP11-181C3.1 | 0.040 | 2.047 |
| FADS1 | 0.033 | 2.045 |
| SLC30A5 | 0.001 | 2.040 |
| CAMK2A | 0.035 | 2.039 |
| UTP11L | 0.003 | 2.035 |
| ZSCAN12 | 0.005 | 2.033 |
| COMMD2 | 0.030 | 2.032 |
| SLC22A5 | 0.002 | 2.024 |
| RP11-466A19.1 | 0.023 | 2.013 |
| BHMT | 0.013 | 2.011 |
| BAIAP2-AS1 | 0.040 | 2.008 |

**Supplemental Table 2.**
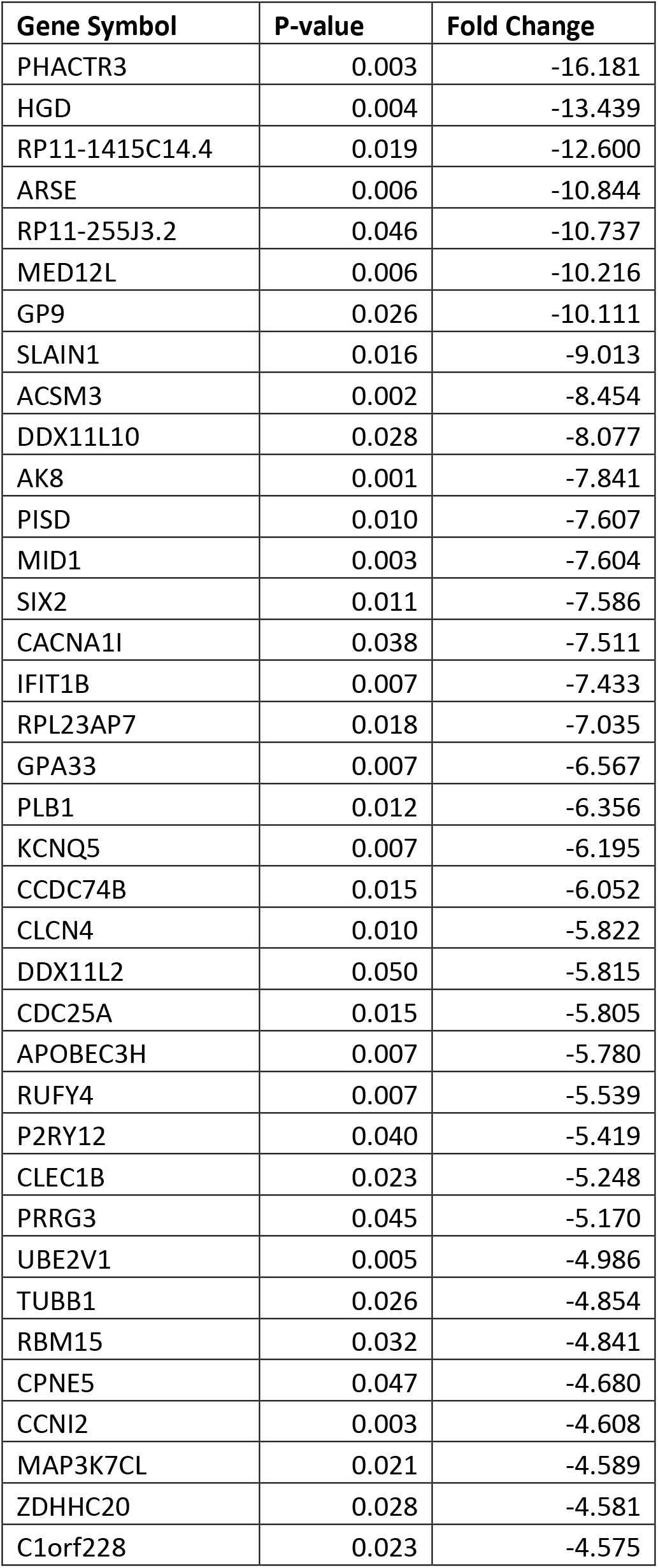
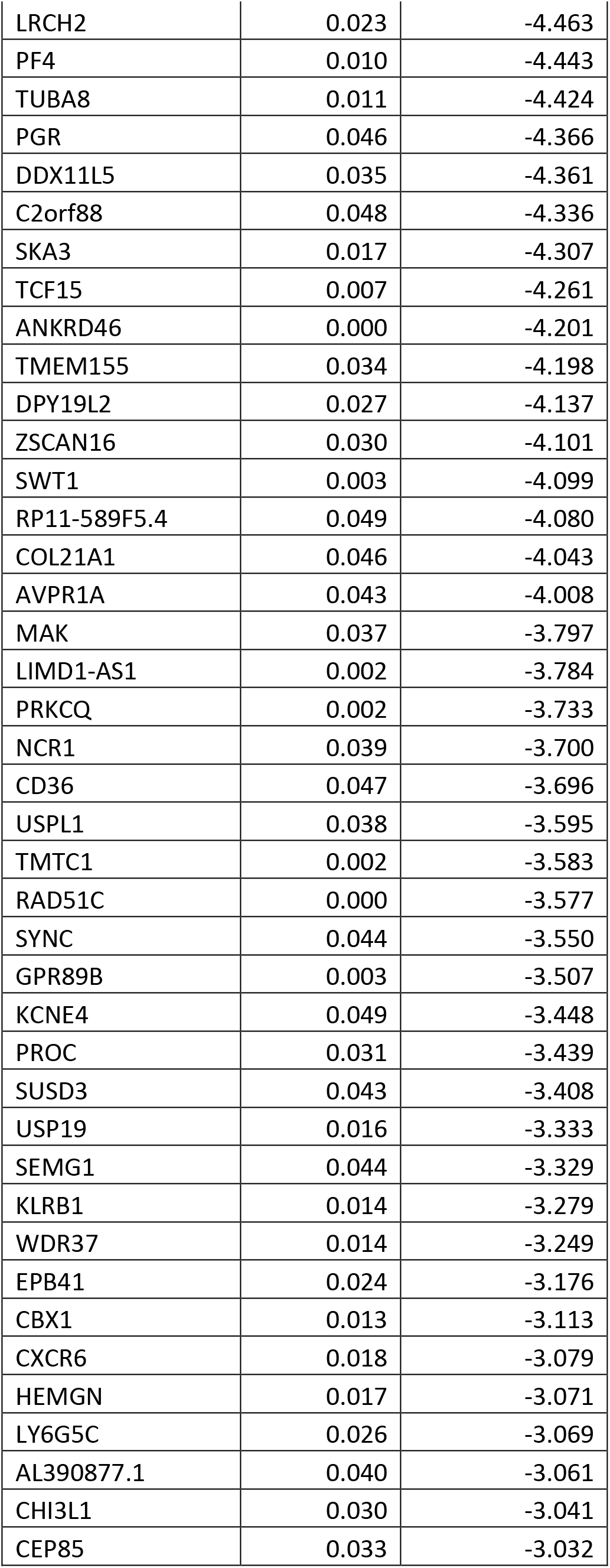
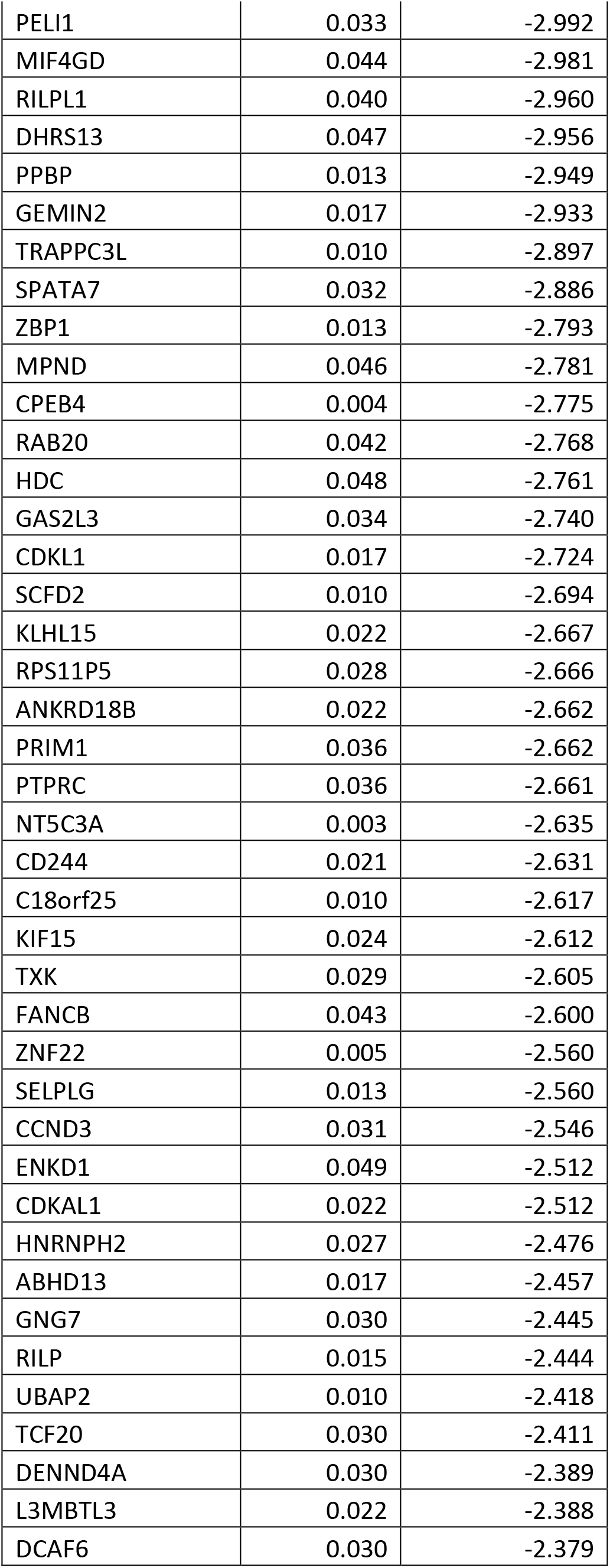
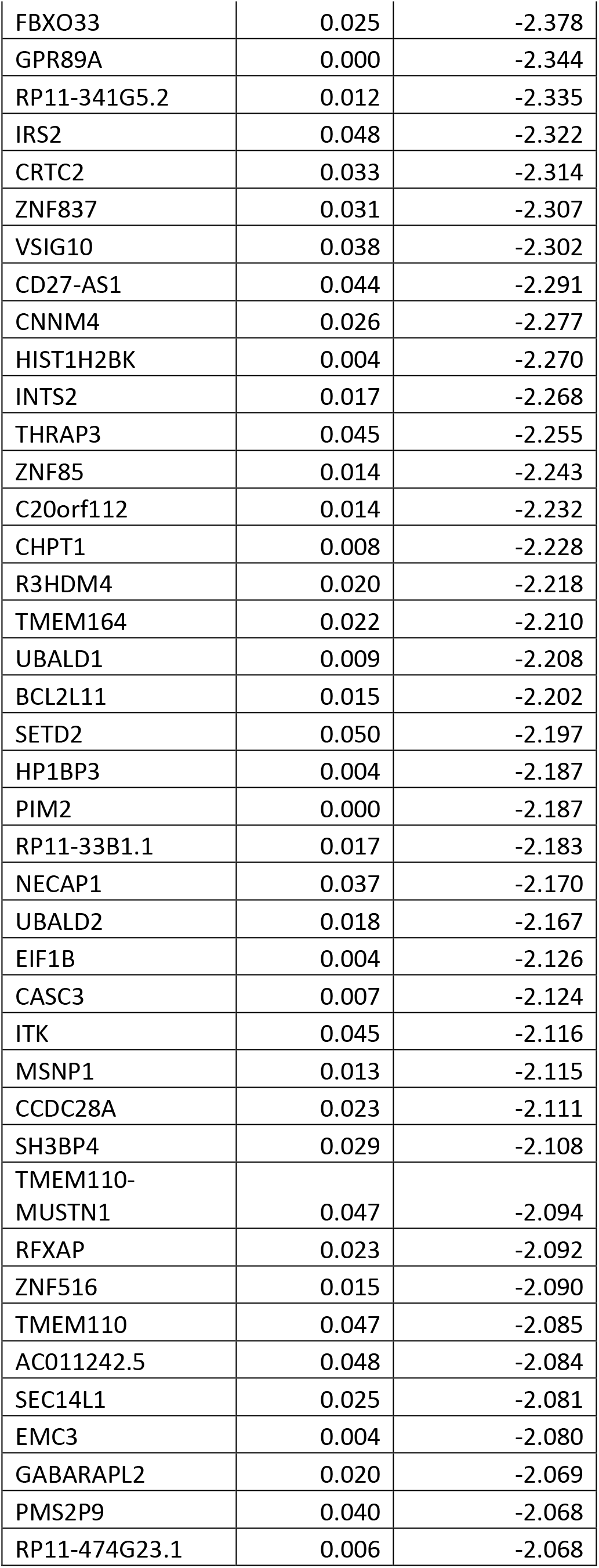
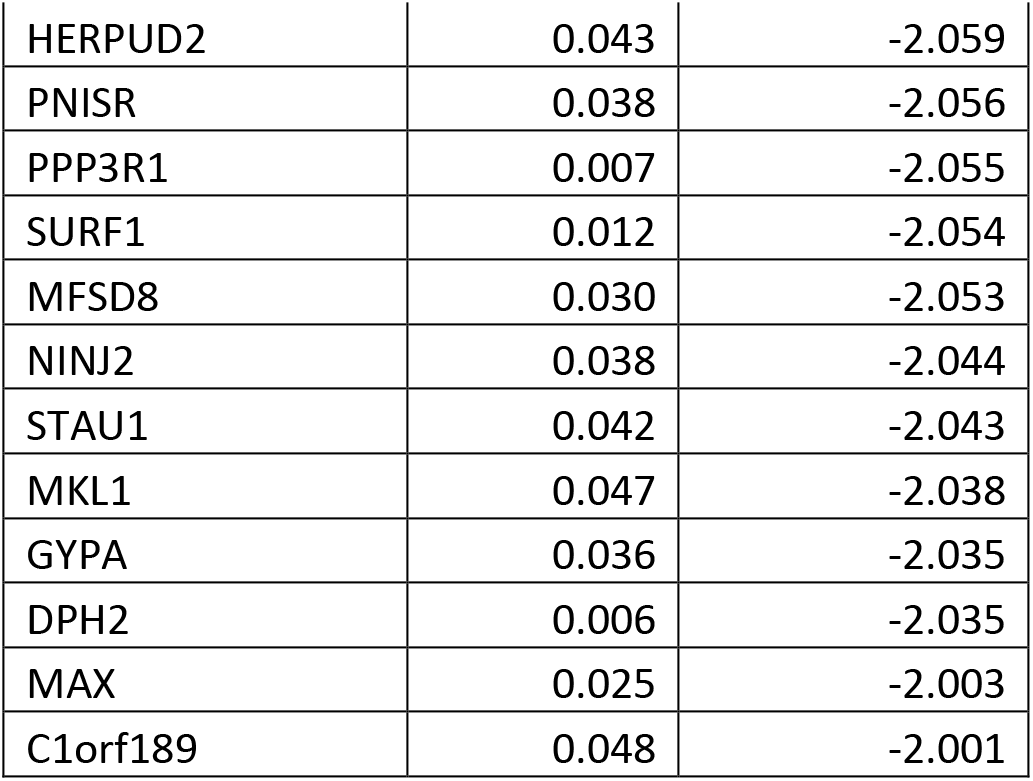
Table of all identified significantly downregulated genes in ultrasound group compared to sham group.

## Notes

### Competing Interest Statement

The authors have declared no competing interest.

## References

1. Fonder, M.A., et al., Treating the chronic wound: A practical approach to the care of nonhealing wounds and wound care dressings. J Am Acad Dermatol, 2008. 58(2): p. 185–206.

2. Ligi, D., L. Croce, and F. Mannello, Chronic Venous Disorders: The Dangerous, the Good, and the Diverse. Int J Mol Sci, 2018. 19(9).

3. Zhao, R., et al., Inflammation in Chronic Wounds. Int J Mol Sci, 2016. 17(12).

4. Nicolaides, A.N., The Most Severe Stage of Chronic Venous Disease: An Update on the Management of Patients with Venous Leg Ulcers. Adv Ther, 2020. 37(Suppl 1): p. 19–24.

5. Valencia, I.C., et al., Chronic venous insufficiency and venous leg ulceration. J Am Acad Dermatol, 2001. 44(3): p. 401–21; quiz 422-4.

6. Franks P., B.J., Collier M., Gethin G., Haesler E., Jawien A., Laeuchli S., Mosti G., Probst S., Weller C., Management of Patients With Venous Leg Ulcers: Challenges and Current Best Practices. Journal of Wound Care, 2016. 25.

7. Weichenthal, M., et al., Low-Frequency ultrasound treatment of chronic venous ulcers. Wound Repair and Regeneration, 1997. 5(1): p. 18–22.

8. Samuels, J.A., et al., Low-frequency (<100 kHz), low-intensity (<100 mW/cm(2)) ultrasound to treat venous ulcers: a human study and in vitro experiments. J Acoust Soc Am, 2013. 134(2): p. 1541–7.

9. Lyu, W., et al., Flexible Ultrasonic Patch for Accelerating Chronic Wound Healing. Adv Healthc Mater, 2021. 10(19): p. e2100785.

10. Ngo O., N.E., Gunasekaran V., Sankar P., Putterman M., Lafontant A., Nadkarni S., DiMaria-Ghalili RA., Neidrauer M., Zubkov L., Weingarten M., Margolis D., Lewin PA., Development of low frequency (20-100 kHz) clinically viable ultrasound applicator for chronic wound treatment. IEEE Transactions Ultrasound Ferroelectric Frequency Control, 2019. 33(3): p. 572–580.

11. Maan, Z.N., et al., Noncontact, low-frequency ultrasound therapy enhances neovascularization and wound healing in diabetic mice. Plast Reconstr Surg, 2014. 134(3): p. 402e–411e.

12. Dunn, L., et al., Murine model of wound healing. J Vis Exp, 2013(75): p. e50265.

13. Grada, A., J. Mervis, and V. Falanga, Research Techniques Made Simple: Animal Models of Wound Healing. J Invest Dermatol, 2018. 138(10): p. 2095–2105 e1.

14. Takao, K. and T. Miyakawa, Genomic responses in mouse models greatly mimic human inflammatory diseases. Proc Natl Acad Sci U S A, 2015. 112(4): p. 1167–72.

15. Nassiri, S., et al., Relative Expression of Proinflammatory and Antiinflammatory Genes Reveals Differences between Healing and Nonhealing Human Chronic Diabetic Foot Ulcers. J Invest Dermatol, 2015. 135(6): p. 1700–1703.

16. Bajpai, A., et al., Effects of Non-thermal, Non-cavitational Ultrasound Exposure on Human Diabetic Ulcer Healing and Inflammatory Gene Expression in a Pilot Study. Ultrasound Med Biol, 2018. 44(9): p. 2043–2049.

17. Yang, Q., et al., Low-Intensity Ultrasound-Induced Anti-inflammatory Effects Are Mediated by Several New Mechanisms Including Gene Induction, Immunosuppressor Cell Promotion, and Enhancement of Exosome Biogenesis and Docking. Front Physiol, 2017. 8: p. 818.

18. Hundt W., Y.E., Steinbach S., Bednarski., Guccione S., Comparison of continuous vs. pulsed focused ultrasound in treated muscle tissue as evaluated by magnetic resonance imaging, histological analysis, and microarray analysis. Molecular Imaging, 2008(18): p. 993–1104.

19. Sindrilaru, A., et al., An unrestrained proinflammatory M1 macrophage population induced by iron impairs wound healing in humans and mice. J Clin Invest, 2011. 121(3): p. 985–97.

